# Triglyceride lipolysis driven by glucose restriction triggers liquid-crystalline phase transitions and proteome remodeling of lipid droplets

**DOI:** 10.1101/2022.04.15.488458

**Authors:** Sean Rogers, Long Gui, Anastasiia Kovalenko, Valeria Zoni, Maxime Carpentier, Kamran Ramji, Kalthoum Ben Mbarek, Amelie Bacle, Patrick Fuchs, Pablo Campomanes, Evan Reetz, Natalie Ortiz Speer, Emma Reynolds, Abdou Rachid Thiam, Stefano Vanni, Daniela Nicastro, W. Mike Henne

**Author notes:** these authors contributed equally to this work (Sean Rogers, Long Gui, Anastasiia Kovalenko, Valeria Zoni).

## Abstract

Lipid droplets (LDs) are reservoirs for triglycerides (TGs) and sterol-esters (SEs), but how these lipids are organized within LDs and influence its proteome remains unclear. Using *in situ* cryoelectron tomography, we show that glucose restriction triggers lipid phase transitions within LDs generating liquid-crystalline lattices inside them. Mechanistically this requires TG lipolysis, which decreases the LD TG:SE ratio, promoting SE transition to a liquid-crystalline phase. Molecular dynamics simulations reveal TG depletion promotes spontaneous TG and SE de-mixing in LDs, additionally altering the lipid packing of the phospholipid monolayer surface. Fluorescence imaging and proteomics further reveal that liquid-crystalline phases are associated with selective remodeling of the LD proteome. Some canonical LD proteins including Erg6 re-localize to the ER network, whereas others remain LD-associated. Model peptide LiveDrop also redistributes from LDs to the ER, suggesting liquid-crystalline-phases influence ER-LD inter organelle transport. Our data suggests glucose restriction drives TG mobilization, which alters the phase properties of LD lipids and selectively remodels the LD proteome.

## Introduction

Lipid droplets (LDs) are unique endoplasmic reticulum (ER)-derived organelles dedicated to the storage of energy-rich neutral lipids. Structurally, LDs are composed of a hydrophobic core of triglycerides (TGs) and sterol-esters (SEs) that is surrounded by a phospholipid monolayer decorated by specific proteins. Beyond their roles in energy homeostasis, recent work highlights the roles of LDs in signaling, development, and metabolism (Welte and Gould, 2017), (Olzmann and Carvalho, 2019), (Walther et al., 2017). These diverse jobs are largely dictated by the LD proteome, but a pervasive question is how specific proteins are targeted to the LD surface. Furthermore, whether the LD proteome is static or dynamic, and how LD lipid composition and cellular metabolic cues influence LD protein residency remains poorly understood.

LDs are generated at the ER and can remain connected to the ER bilayer for extended periods (Jacquier et al., 2011), (Kassan et al., 2013). As such, Type I LD proteins can translocate between the ER and LD monolayer via lipidic bridges connecting the two organelles (Wilfling et al., 2013). Elegant *in vitro* studies have suggested that LD protein targeting promotes energetically favorable conformational changes within some proteins, and the re-positioning of proteins to LDs from the ER network can even influence their enzymatic activities, or modulate their degradation (Caillon et al., 2020), (Chorlay and Thiam, 2020), (Leber et al., 1998), (Schmidt et al., 2013), (Ohsaki et al., 2006). A second mechanism of LD targeting occurs from the cytoplasm, where soluble proteins insert into the LD monolayer via a hydrophobic region, amphipathic helix, or lipid moiety. Here hydrophobic protein regions recognize lipid packing defects (LPDs) between the phospholipid monolayer lipid head groups, enabling their insertion into the neutral lipid core (Bacle et al., 2017), (Prevost et al., 2018), (Chorlay and Thiam, 2020), (Chorlay et al., 2021).

Although monolayer phospholipids can regulate LD protein targeting, how neutral lipids (NLs) within the LD core influence protein localization is less understood. However, NLs clearly impact the composition of the LD surface proteome (Chorlay and Thiam, 2020). For example, in yeast some proteins preferentially decorate TG-rich LDs (Gao et al., 2017). Molecular studies also indicate that protein insertion into the LD neutral lipid core enables proteins to fold with lower free energy, and polar residues within hydrophobic regions can even interact with TG, further anchoring them to the LD (Olarte et al., 2020). Despite these insights, how NL pools ultimately influence the composition and dynamics of the LD proteome is underexplored, yet central to our understanding of LD organization and functional diversity.

NLs generally form an amorphous mixture within the hydrophobic LD core. This organization, and even the phase properties of LD lipids themselves, can change in response to various cellular stimuli. HeLa cells induced into mitotic arrest or starvation exhibit lipid phase transitions within their LDs, generating liquid-crystalline lattices (LCLs) inside the LD core that can be visualized via cryo-electron tomography (cryo-ET) as striking onion-like layers inside the LDs (Mahamid et al., 2019). Yeast biochemical studies also proposed similar segregation of TGs and SEs into discrete layers within LDs (Czabany et al., 2008). This lipid reorganization is attributed to the biophysical properties of SEs, which can transition from disordered to ordered smectic phases under physiological conditions (Kroon, 1981), (Ginsburg et al., 1984), (Shimobayashi S, 2019), (Czabany et al., 2008). Such phase transitions are also associated with human pathologies including atherosclerosis, and liquid-crystalline LDs were even observed in the macrophage of a patient with Tangier disease (Lundberg, 1985), (Katz et al., 1977). How these phase transitions are triggered, and whether they influence organelle physiology, or are simply a biophysical consequence of the properties of SEs, is unknown.

Here, we utilized budding yeast to dissect the metabolic cues governing lipid phase transitions within LDs. We used cryo-ET of cryo-focused ion beam (cryo-FIB) milled yeast cells to study the *in situ* architecture of LDs in their native environment, under ambient or glucose-restricted conditions. We show that in response to acute glucose restriction (AGR), yeast initiate TG lipolysis, which induces the formation of LCLs within LDs. The formation of LCL-LDs closely correlates with changes in the TG:SE lipid ratios within the LD core. In line with this, molecular dynamics (MD) simulations of model LDs indicate that variations in this TG:SE ratio are sufficient to promote their spontaneous de-mixing within LDs. This lipid de-mixing also alters the LD monolayer surface, and consistently we find that LCL-LD formation drastically alters LD protein targeting and selectively changes the LD surface proteome. Taken together, our data suggest that upon glucose restriction, TG lipolysis triggers spontaneous lipid demixing within LDs that supports liquid-crystalline phase transitions and the subsequent remodeling of the LD surface proteome.

## Results

### Acute glucose restriction promotes TG lipolysis-dependent liquid-crystalline phase transitions in LDs

Previous studies from our group indicated that budding yeast exposed to acute glucose restriction (AGR), where yeast are transferred from a glucose-rich (2%) synthetic complete media to a low-glucose (0.001%) media, exhibit metabolic remodeling that favors the production of SEs, which are stored in LDs (Rogers et al., 2021). We used cryo-ET to investigate if AGR also impacts LD morphology. We rapidly froze yeast cells that were either in logarithmic (logphase) growth in glucose-rich media, or exposed to 4 hrs of AGR, and used cryo-FIB milling to generate 100-200-nm-thick lamellae of the vitrified cells. These lamellae were then imaged by cryo-ET to reveal the three-dimensional (3D) structure of native LDs *in situ.* The cryo-FIB milled lamella exhibited a well-preserved yeast ultrastructure, including the nucleus, vacuole, mitochondria and LDs (**Figure 1A, Figure S1, SMovie 1**). Typical LDs could be distinguished from other cellular organelles by their relatively electron-dense, amorphous interior that was surrounded by a thin phospholipid monolayer (**Figure 1B, SMovie 2**). In contrast to normal LDs in glucose-fed log-phase cells, ~77% of the LDs observed in 4hrs AGR-treated yeast displayed reorganization of their interior, including the appearance of distinct concentric rings in the LD periphery (**Figure 1 C-D, M for quantification, SMovie 3**). These rings appear similar to lattices previously observed in liquid-crystalline-phase LDs, which exhibited a regular spacing of ~3.4-3.6nm between their layers, suggesting they were composed of sterol-esters (SEs) (Mahamid et al., 2019), (Engelman and Hillman, 1976). Indeed, our line-scan analysis showed regular 3.4nm spacing between rings (**Figure 1E**), suggesting these LDs indeed exhibited SE liquid-crystalline lattices (LCLs). Thus, we refer these “onion-like” LDs as LCL-LDs. Notably, these were never observed in the log-phase yeast (**Figure 1B, M**).

**Figure 1:**
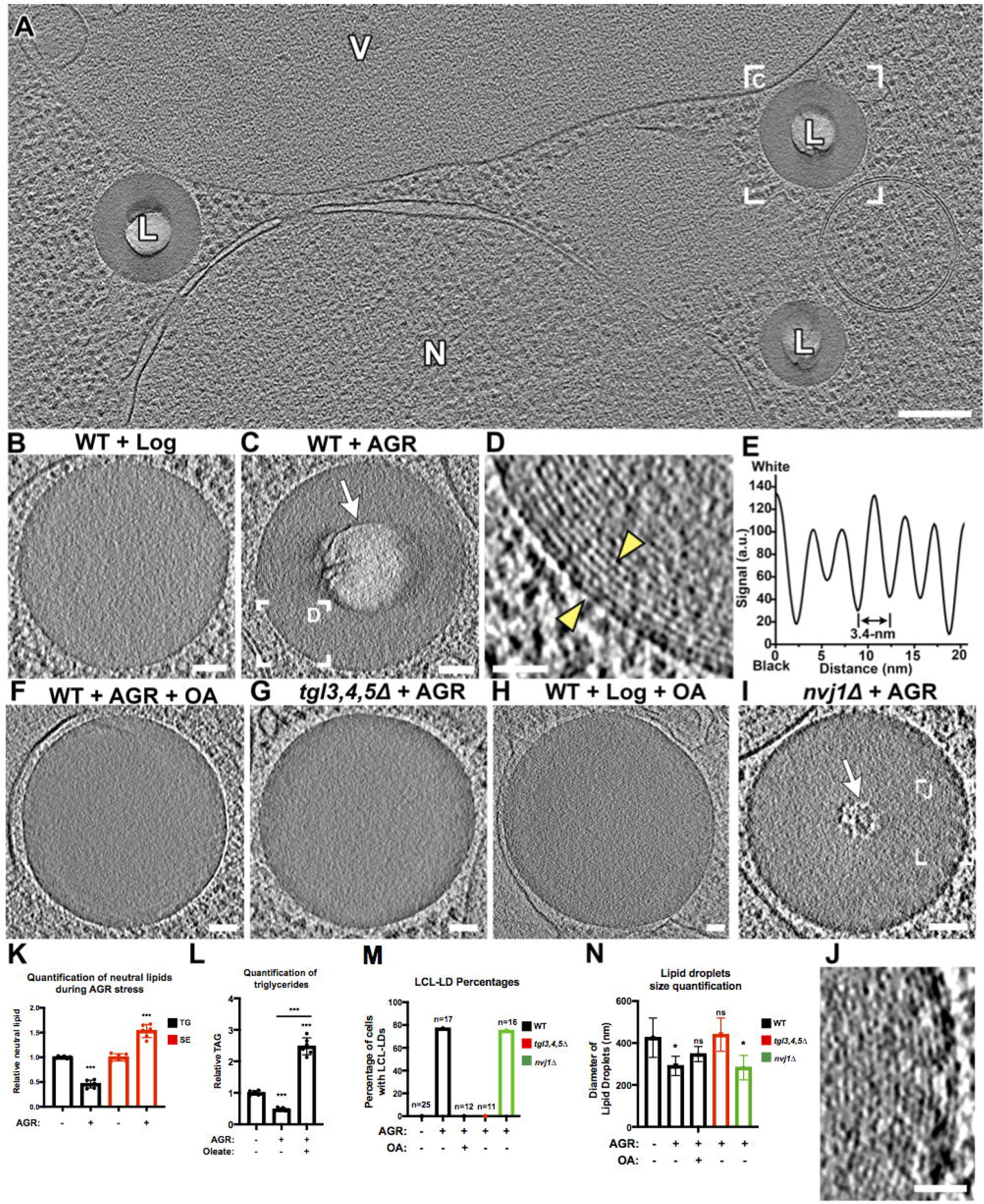
Visualization of the liquid-crystalline layers in lipid droplets (LCL-LD) promoted by TG lipolysis using *in situ* cryo-ET. **A**) Representative tomographic slice from a cryo-FIB-milled and cryo-ET reconstructed wildtype (WT) yeast cell grown for 4hrs under acute glucose restriction (AGR). Note the “bubbled” (lighter) centers of the LDs (L). V, vacuole. N, nucleus. A different tomographic slice of the boxed LD is also shown in (**C**). **B-J**) Representative tomographic slices of LDs in yeast from glucose-fed WT in log phase (**B**), WT after 4hrs AGR (**C**, boxed area magnified in **D, E** shows line-scan plot of area between yellow arrowheads), WT after 4hrs AGR + 0.1% oleate (OA) (**F**), *tgl3,4,5Δ* yeast after 4hrs AGR (**G**), WT cultured with 2% glucose and 0.1% OA (**H**), *nvj1Δ* after 4hrs AGR (**I**, and boxed area magnified in **J**). Liquidcrystalline layers (LCL) were only observed in LDs from WT and *nvj1Δ* yeasts in AGR (**C**, **D**, **I**, **J**). White arrows highlight the ‘bubbles’ due to electron radiation in centers of LCL-LDs. **K**) Quantification of relative whole-cell TGs and SEs in log and 4hrs AGR conditions. **L**) Relative TGs in log and 4hrs AGR conditions. **M, N**) % abundance of LCL-LDs (**M**) and diameters of LDs (**N**) under various conditions measured in cryo-tomograms. Note that the observed diameter depends on the plane at which the LDs were sectioned; therefore, for size measurements, only LDs with clearly visible monolayer (indicating a slice through the LD center) were included. Scale bars: 200nm (A). 50nm (B-C, F-I). 20nm (D, J).

In addition to the LCLs beneath the LD monolayer surface, the amorphous centers of LCL-LDs were unusually sensitive to electron radiation, causing excessive radiolysis and “bubbling” (i.e. the generation of a gas bubble trapped in the ice that appears white in cryo-EM images) during tilt-series acquisition (**Figure 1C, white arrow**). This increased radiation sensitivity was only observed in LCL-LDs, but not in LDs with entirely amorphous interiors (i.e. not observed in the 23% disordered LDs of AGR-treated yeast, nor in any LDs of log-phase yeast). We generated comparative ‘bubblegrams’, (i.e. a series of 2D cryo-EM images where the same sample area was exposed to an increasing amount of electron dose), which revealed that the centers of LCL-LDs exhibited bubbling following exposure to <30 e/Å^2^, whereas amorphous LDs from log-phase yeast did not show any bubbling even at 400 e/Å^2^ dosages (**Figure S1 A-J**). Previous studies of electron radiation-induced bubbling of frozen biomolecules in aqueous solution and within cells demonstrated that similar gas bubbles contained mostly molecular hydrogen gas which became trapped in the vitrified samples (Leapman and Sun, 1995), (Aronova et al., 2011). Although the mechanism of radiation-induced bubbling and increased radiation-sensitivity within the center of LCL-LDs is not clear, it may be due to the production of gases derived from a specific combination of lipids or metabolites present within LCL-LDs.

We hypothesized that the formation of LCL-LDs may involve changes to the NLs within the LD core. To investigate the effects of AGR on yeast NL pools, we monitored TG and SE levels in log-phase and 4hrs AGR-treated yeast. Indeed, whereas log-phase yeast contained similar levels of TG and SE, AGR treated yeast contained significantly less TG (**Figure 1K)**. As expected, AGR yeast also had increased amounts of SEs (**Figure 1K)**, as previously observed (Rogers et al., 2021), indicating the TG:SE ratio within the LDs was significantly decreased to ~0.5:1.5 compared to a normal ratio of ~1:1 (Leber et al., 1994). We hypothesized that LCL-LD formation was promoted by TG loss from LDs. To test this, cryo-ET was performed on yeast lacking the major TG lipases *(tgl3,4,5Δ).* Indeed, 4hrs AGR treated *tgl3,4,5Δ* yeast did not form any detectable LCL-LDs (**Figure 1G, M**), suggesting TG lipolysis was required for LCL-LD formation. In support of this, LDs in wildtype (WT) AGR-treated yeast were significantly smaller in diameter than log-phase LDs, and this reduced size was suppressed in *tgl3,4,5Δ* yeast (**Figure 1N**), suggesting the size reduction was due to lipid loss via TG lipolysis.

To further dissect how TGs influence LCL-LDs, we treated yeast with 0.1% oleic acid (OA), which promotes TG synthesis. As expected, OA elevated cellular TG levels in yeast when cultured in its presence during the 4hrs AGR treatment (**Figure 1L**), and notably no LCL-LDs were observed during log-phase nor in this AGR+OA condition (**Figure 1F, H, M**). In line with this, whereas LD sizes in AGR-treated yeast were significantly smaller than in log-phase cells, their sizes slightly recovered under the AGR+OA condition **(Figure 1F, N)**. Since we previously observed that the yeast nucleus-vacuole junction (NVJ) can serve as a site for LD biogenesis during nutrient stress (Hariri et al., 2018), we also examined whether NVJ loss impacted LCLLD formation. Cryo-ET of *nvj1Δ* yeast cells showed the expected loss of tight contacts between the outer nuclear envelope and the vacuole **(Figure S1K, L)**. However, *nvj1Δ* yeast exhibited ~75% LCL-LDs under AGR conditions (very similar to WT yeast), indicating that the NVJ was not required for LCL-LD formation (**Figure 1I, J, M**).

Since SEs can form liquid-crystalline lattices, we next tested whether SEs were required for LCL-LD formation. We monitored LDs in *are1are2Δ* yeast that cannot synthesize SEs. Surprisingly, in 15 different cryo-FIB lamella of *are1are2Δ* yeast cells, no LDs could be observed (**Figure S1M**). However, fluorescence staining with monodansylpentane (MDH) LD stain confirmed the presence of LDs in *are1are2Δ* yeast during AGR stress, but these LDs were small and sparser compared to any of the other examined strains (**Figure S1N**). Thus, the reduction in LD size and abundance within cells may account for the inability to observe LDs in the cryotomograms of the 100-200nm thick lamellae.

Collectively, these data suggest that TG abundance is a key modulator of SE-associated phase transitions within LDs. They support a model where Tgl-dependent TG lipolysis during AGR promotes LCL-LD formation by depleting the TG pool that maintains SE in its disordered phase.

### TG and SE spontaneously separate in model LDs in response to altered TG:SE ratios

To investigate the mechanism by which AGR treatment promotes the formation of LCL-LDs, we performed MD simulations of model LDs, as this technique has been shown to provide molecular information on LD-like lipid assemblies (Ben M’barek et al., 2017), (Bacle et al., 2017), (Zoni et al., 2021), (Prevost et al., 2018), (Chorlay et al., 2019). Since cryo-ET suggests that during AGR treatment SEs are enriched at the LD periphery (**Figure 1**), and biochemical measurements indicate that this phenomenon correlates with imbalanced TG:SE ratios within LDs (**Figure 1K**), we initially performed MD simulations to investigate how SEs and TGs partition within the LD core at different ratios. We built lipid trilayers, model systems in which different concentrations of SEs and TGs are sandwiched between two monolayers of phosphatidylcholine (PC) phospholipids, representing the LD surface, and surrounded by water (**Figure 2A**). These structures mimic mature LDs, and they were previously used to study LD structural properties (Bacle et al., 2017) and protein targeting to LDs (Prevost et al., 2018).

**Figure 2:**
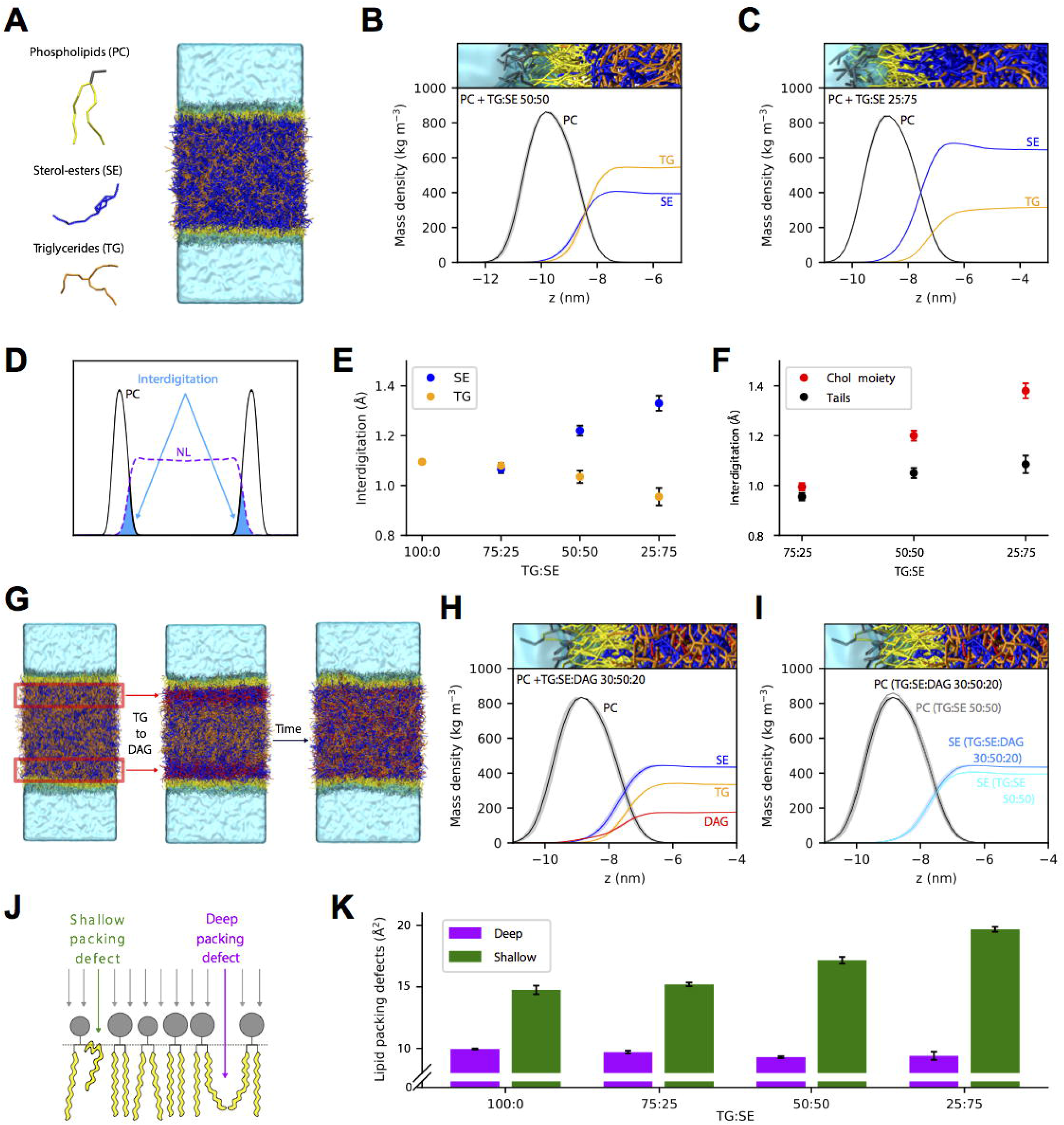
SE and TG partition non-uniformly in the core of *in silico* model LDs. **A**) Trilayer system of molecular dynamics (MD) simulations, composed of phosphatidylcholine (PC) phospholipids, yellow and gray, sterol-esters (SE), blue, and triglycerides (TG), orange. **B**) Density profile of TG, SE and PC in a trilayer system containing 50:50 TG:SE. Only a small region of the profile is shown for clarity. For density profiles, the results are reported as the average between the analysis of the first part and the last part of the simulations, while the shaded regions correspond to the area between the two curves. See Methods for details. **C**) Density profile of TG, SE and PC in a trilayer system containing 25:75 TG:SE. **D**) Schematic representation of interdigitation and calculations for systems at different TG:SE ratios. The results are reported as the average between the analysis of the first part and the last part of the simulations, while the error bars correspond to the distance between the results. See methods for details. **E**) Interdigitation for SE and TG at different TG:SE ratios. **F**) Interdigitation between PC lipids and the cholesterol moiety (red) or the acyl chain (black) of SE molecules at different TG:SE ratios. **G**) Representative snapshots describing the time evolution of simulations mimicking TG lipolysis into DAG. **H)** Density profiles of PC, SE, TG and DAG after equilibration. **I)** Comparison of PC and SE density profile for trilayer containing 50% SE in the absence (cyan) or presence (light blue) of 30% DAG. **J**) Schematic representation of shallow and deep lipid packing defects (LPDs). **K**) Quantification of deep and shallow lipid packing defects (LPDs) in systems with different TG:SE ratios.

As a reference starting point, we investigated a model LD trilayer with a NL ratio of 50% SE and 50% TG molecules (TG:SE 50:50), as this composition is compatible with that of WT yeast cells in log-phase (**Figure 1K**). Starting from an initial random distribution of SEs and TGs in the trilayer core, the simulations show that when equilibrium is reached, SEs and TGs are similarly enriched close to the PC monolayers (**Figure 2B, Figure S2A**). Of note, this comparable behaviour of TGs and SEs is quite unexpected, since in oil-water interfaces of TG:SE mixtures in the absence of phospholipids, TG molecules are acutely enriched at the oil-water interface **(Figure S2B)**, because of the lower interfacial tension of TGs with water in comparison with that of SEs (**STAR Figure 1B**). Thus, our data suggest that preferential interactions with PLs drive SE molecules close to the LD surface.

To mimic the variations in NL composition upon AGR treatment, we next modelled the LD core at a higher (25:75 TG:SE) SE concentration (**Figure 2C, Figure S2C**) and compared it with the previous (50:50 TG:SE) composition. Remarkably, even in the absence of any transition to the LCL state, in this condition there is a clear difference in surface localization of TG versus SE, with SE molecules becoming more abundant at the LD surface at the higher SE concentration (**Figure 2C, Figure S2C**). Of note, this NL composition corresponds approximately to that of 4hrs AGR-treated yeast, where LCL-LDs were observed (**Figure 1C,K**).

As an additional measure of the extent to which NLs pervade the LD interfacial region with water, that is mostly populated by phospholipids, we next measured the interdigitation between NLs and the PC monolayer (**Figure 2D**). Remarkably, simulations show that as the SE relative concentration with respect to TG increases (TG:SE ratios of 75:25 → 50:50 → 25:75), SEs interdigitate more with the phospholipid surface than TGs (**Figure 2E**). Since the interdigitation between the surface PC monolayer and the acyl chain of SE molecules remains approximately constant for increasing SE concentrations, our simulations suggest that this mechanism may be driven by the preferential interaction of the cholesterol moiety of SE with surface phospholipids (**Figure 2F**). Taken together, our results suggest that phospholipids preferentially interact with SEs, leading to a non-uniform distribution of the two NLs in the LD core. This non-uniform partitioning is broadened at TG:SE ratios where SE is in abundance, such as those observed under yeast AGR treatment.

Since experiments suggest that the action of TG lipases is necessary for LCL-LD transition (**Figure 1G,M**), we wondered how the products of TG lipolysis would partition in the LD core. Assuming that fatty acids released from TG lipolysis would be directly transported away from LDs for cellular bioenergetics, we tested how the other main product of TG metabolism, diacylglycerol (DAG), distributes in the LD core. To mimic a physiological situation where DAGs are produced at the LD periphery where TG lipases act, we modified only the TG molecules that are close to the phospholipid monolayers to DAGs, and then monitored their spatial localization over time (**Figure 2G**). At equilibrium, DAGs partitioned similarly to TGs near the centre of the LD interior, with SEs again being the predominant lipid species at the LD periphery, immediately below the phospholipid surface (**Figure 2H, Figure S2D)**. A direct comparison of SE density in the presence/absence of DAG molecules indicates that even in the presence of more surface-active TG products, such as DAG, SEs retain their surface localization (**Figure 2I)** (Campomanes et al., 2019), suggesting that TG lipolysis induced by TG lipases is unlikely to alter the surface enrichment of SE molecules.

Collectively, our simulations support the hypothesis that, at high concentrations of SEs within the LD core, SEs are enriched near the phospholipid monolayer surface. This suggests that the non-uniform partitioning of TG and SE molecules at the LD periphery could be attributed to physicochemical properties of these molecules already in the liquid state, *i.e.* when the LD core is amorphous. Of note, TGs are not completely excluded from the LD periphery in this modelling, suggesting that, even at high SE concentrations, TGs are potentially still accessible to surface lipases. These observations could explain why, even in our simulations at 25:75 TG:SE ratio, we do not observe crystalline lattice formations. It is possible TGs need to be almost completely removed from the LD periphery to promote the phase transition to LCL-LDs.

### Spontaneous NL separation alters the LD monolayer surface properties

Since SEs and TGs can distribute non-homogenously within the LD core, we next investigated whether this de-mixing influences the surface properties of LDs. We computed lipid packing defects (LPDs), a measure of the stable voids between the phospholipid headgroups that expose lipid hydrophobic regions to water. LPDs can be classified as “deep” if they extend below the phospholipid glycerol backbone, or “shallow” if they are only above the phospholipid glycerol backbone (**Figure 2J**), and both types have been identified as crucial to modulate peripheral protein binding to bilayers (Vanni et al., 2014), (Vanni et al., 2013) and monolayers (Prevost et al., 2018), (Olarte et al., 2020), (Caillon et al., 2020), (Chorlay and Thiam, 2020). Our simulations show that the number of deep LPDs remains almost constant in systems with increasing concentrations of SEs (**Figure 2K**). However, we observed a significant increase in the number of shallow LPDs when SEs rose above 50% in the trilayer core (**Figure 2K**). This may be due to the increased interdigitation of the cholesterol moiety of SEs, which we found was more prone to interdigitate in the acyl chain layer of the PC phospholipid surface of the trilayer system at higher SE concentrations (**Figure 2F**).

In summary, our MD simulations support a model where SEs and TGs can spontaneously demix at the surface of LDs when SEs are in excess. Such conditions closely match the empirically measured NL abundances we observe in yeast during AGR, where we observe the appearance of LCL-LDs in a TG lipolysis-dependent manner. However, a limitation of our simulations is that they do not directly model the liquid-crystalline phase of the SEs. Even if these simulations do not exhibit phase transitions at high SE concentrations, the observations support a model where local SE enrichment near the LD monolayer surface can alter the monolayer properties, and thus may influence LD protein targeting. It may also explain why earlier studies on amorphous LDs show a different proteome targeting SE-rich vs TG-rich LDs (Khor et al., 2014), (Hsieh et al., 2012), (Gao et al., 2017).

### LCL-LD formation correlates with Erg6 re-distribution from LDs to the ER network

Motivated by our *in silico* data indicating that variations in TG:SE ratios can alter LD monolayer surface properties, we next investigated whether AGR stress and its associated LCL-LD formation impact LD protein targeting. Indeed, changes in the LD surface monolayer have been proposed to control protein targeting to lipid bilayers (Vanni et al., 2014), (Vanni et al., 2013) and monolayers (Prevost et al., 2018), (Caillon et al., 2020), (Chorlay and Thiam, 2020), (Olarte et al., 2020). This observation is also in agreement with previous studies indicating that LD proteins may interact with TGs contained within the LD interior (Olarte et al., 2020), (Santinho et al., 2021). However, whether smectic LCL lipid phase transitions influence LD protein targeting remains unknown. To explore this possibility, we imaged the canonical yeast LD protein Erg6 tagged with mNeonGreen (Erg6-mNg) over time in yeast exposed to AGR stress. As expected, Erg6-mNg initially colocalized with LDs prior to AGR stress (t=0). However, the Erg6-mNg labeling pattern changed after ~1hr AGR, and primarily decorated the cortical ER and nuclear envelope (**Figure 3A**). Erg6-mNg remained at the ER network throughout 2, 4, and 24 hrs AGR, and notably the LD stain gradually dimmed over these time-points, consistent with the loss of LD volume via lipolysis.

**Figure 3:**
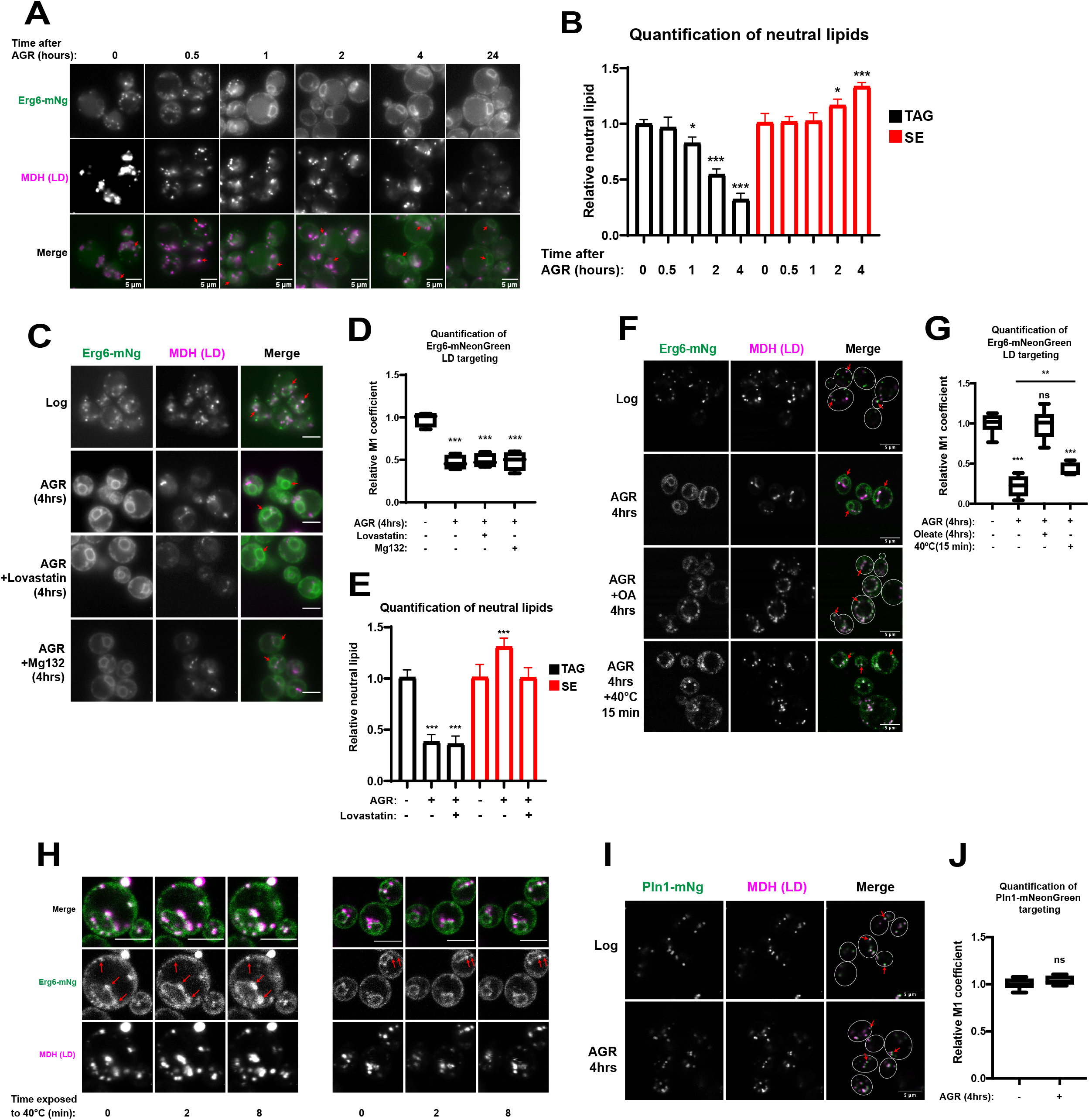
Erg6 LD de-localization correlates with altered TG:SE ratios and LCL-LD formation. **A**) Yeast expressing Erg6-mNeonGreen (mNg) and stained for LDs (monodansylpentane, MDH) at time-points when yeasts were transferred from log-phase (2% glucose) to acute glucose restriction (AGR). Red arrows indicate protein targeting. **B**) TLC measurements of TG and SE in AGR-treated yeast over time. **C**) Yeast expressing Erg6-mNg and MDH-stained for LDs. Yeast were either treated with AGR, AGR+lovastatin (sterol synthesis inhibitor) together, or AGR+Mg132 (proteasome inhibitor) together for 4 hrs. **D**) Relative Manders M1 coefficient of Erg6-mNg colocalization with MDH LD stain in various conditions related to C. **E**) TLC measurements of TG and SE from yeast treated for 4 hrs with AGR or AGR+lovastatin. **F**) Yeast expressing Erg6-mNg and MDH-stained for LDs in log-phase (2% glucose), AGR, and AGR+0.1% oleate (OA), and AGR+15min 40°C. **G**) Relative Manders M1 coefficient of Erg6-mNg colocalization with LD stain MDH in various conditions from F. **H**) Timelapse imaging of Erg6-mNg yeast stained with MDH, and heated up to 40°C following 4hrs AGR**. I**) Pln1(Pet10)-mNg in log and 4hrs AGR. **J**) Relative M1 coefficient of Pln1-mNG with LD targeting. Statistics are one-way ANOVA. Scale bars 5μm.

Using thin layer chromatography, we also observed a gradual decrease in cellular TG during the AGR time-course, with approximately 20 and 40 percent reduction in TG after 1 and 2 hrs AGR, respectively (**Figure 3B)**. As previously demonstrated, cellular SEs increased during this AGR time-course. However, notably Erg6-mNg delocalization from LDs preceded this SE increase, and this SE increase occurred after the initiation of TG depletion. This suggested that TG depletion, rather than SE increase, may be more important in promoting Erg6-mNg signal loss of LDs and accumulation at the ER. Consistent with this, treating cells with lovastatin, a sterol pathway inhibitor that blunted the SE increases in AGR but did not alter the TG decrease, did not prevent Erg6-mNg accumulation in the ER (**Figure 3C, E**). This collectively suggests that TG depletion, rather than SE elevation, strongly correlates with loss of Erg6-mNg LD signal and its accumulation at the ER network during the AGR time-course.

To rule out the possibility that Erg6-mNg appearance at the ER during AGR was caused by selective turnover of LD-localized Erg6-mNg followed by new synthesis of Erg6 at the ER network, we treated cells with the proteasome inhibitor Mg132 during AGR. Mg132-treated cells still displayed ER localized Erg6-mNg during AGR together with a corresponding decrease of LD-associated Erg6-mNg signal, suggesting the loss of LD-associated Erg6-mNg was not due to proteasomal turnover (**Figure 3C**). To quantify all this, we calculated the Manders M1 coefficient for each condition, which represents the amount of Erg6-mNg signal that overlaps with the LD stain MDH (**Figure 3D**). Indeed following 4 hrs of AGR stress, LD-associated Erg6-mNg signal was reduced by ~50 percent, and this was unaffected upon addition of either lovastatin or Mg132. Collectively, these data suggest that neither SE synthesis nor proteasomal degradation play a significant role in the re-distribution of Erg6-mNg during AGR, and support a model where TG lipolysis promotes Erg6-mNg re-distribution from LDs to the ER network.

Because loss of Erg6-mNg from LDs closely correlated with conditions that promoted LCL-LD formation, we next assessed whether Erg6-mNg maintained LD targeting in conditions that prevented or reversed LCL-LD formation in cryo-ET. First, we monitored Erg6-mNg localization in cells subjected to AGR stress in the presence of 0.1% OA, or upon genetic ablation of TG lipases, both of which suppressed LCL-LD formation. Indeed, Erg6-mNg remained on LDs in both these conditions, suggesting that Erg6-mNg de-localization from LDs tightly correlates with LCL-LD formation and its associated TG reduction (**Figure 3F, G, Figure S3A**). To more directly test whether the biophysical properties of LCL-LDs influenced Erg6-mNg localization, rather than other metabolic changes attributed to AGR stress, we briefly heated Erg6-mNg expressing yeast after 4hrs AGR to 40°C, which is above the predicted phase transition temperature for smectic-phase SEs. Indeed, Erg6-mNg significantly, although not fully, relocalized from the ER network to LDs after only 15 minutes at 40°C (**Figure 3F, G**). We also conducted live-cell imaging of AGR-treated Erg6-mNg yeast during this heating, which revealed a rapid re-distribution of Erg6-mNg from the ER network to LDs within ~8 minutes of applied 40°C heating (**Figure 3H**). Together, these observations support a model in which TG lipolysis and the associated LCL-LD formation promotes Erg6-mNg re-localization from LDs to the ER network, but this targeting can be quickly reversed at higher temperatures, enabling Erg6-mNg to return to the LD surface.

Next, we investigated whether AGR caused a general re-localization of other canonical LD proteins. Surprisingly, Pln1-mNg, a perilipin-like protein also known as Pet10 (Gao et al., 2017), was still detectably localized to LDs by imaging following 4hrs of AGR, suggesting the delocalization of LD proteins during AGR and its associated LCL-LD formation may be selective (**Figure 3I, J**).

### Erg6 and Pln1 LD targeting is influenced by LD neutral lipid content in human cells independent of AGR stress

Since our observations indicated that Erg6 could reversibly target between LDs and the ER network during AGR-associated LCL-LD formation, we next wanted to dissect this dynamic relocalization in an orthogonal cellular system that was uncoupled from the metabolic changes associated with yeast AGR stress. To this end, we expressed eGFP-tagged Erg6 in human HeLa cells, then treated them either for 24hrs with OA to induce TG-rich LDs, or with cholesterol coupled to methyl-β-cyclodextrin to induce the formation of SE-rich LDs (**Figure 4A**). Indeed, cholesterol-treated HeLa cells contained LDs with anisotropic birefringent properties when illuminated with polarized light, consistent with the formation of liquid-crystalline cholesterol-esters within these LDs (Shimobayashi S, 2019) (**Figure 4A,C**). In contrast, the OA-treated TG-rich HeLa cells contained LDs that were not birefringent. Erg6-eGFP was detected on nearly all LDs in both TG-rich and SE-rich LD conditions, but appeared slightly dimmer on LDs in the SE-rich samples, where a dim ER network Erg6-eGFP signal was also observed (**Figure 4C**). Consistent with this, when we calculated the Erg6-eGFP LD-to-ER ratio for each condition, we found they significantly differed. In cells containing TG-rich LDs, Erg6-eGFP exhibited a significantly higher LD-to-ER ratio compared to Erg6-eGFP in the cholesterol-treated cells (**Figure 4C,E**). This suggested that, similar to the observations in yeast, Erg6 exhibited more targeting to TG-rich LDs as compared to SE-rich LDs (**Figure 3A-G**).

**Figure 4:**
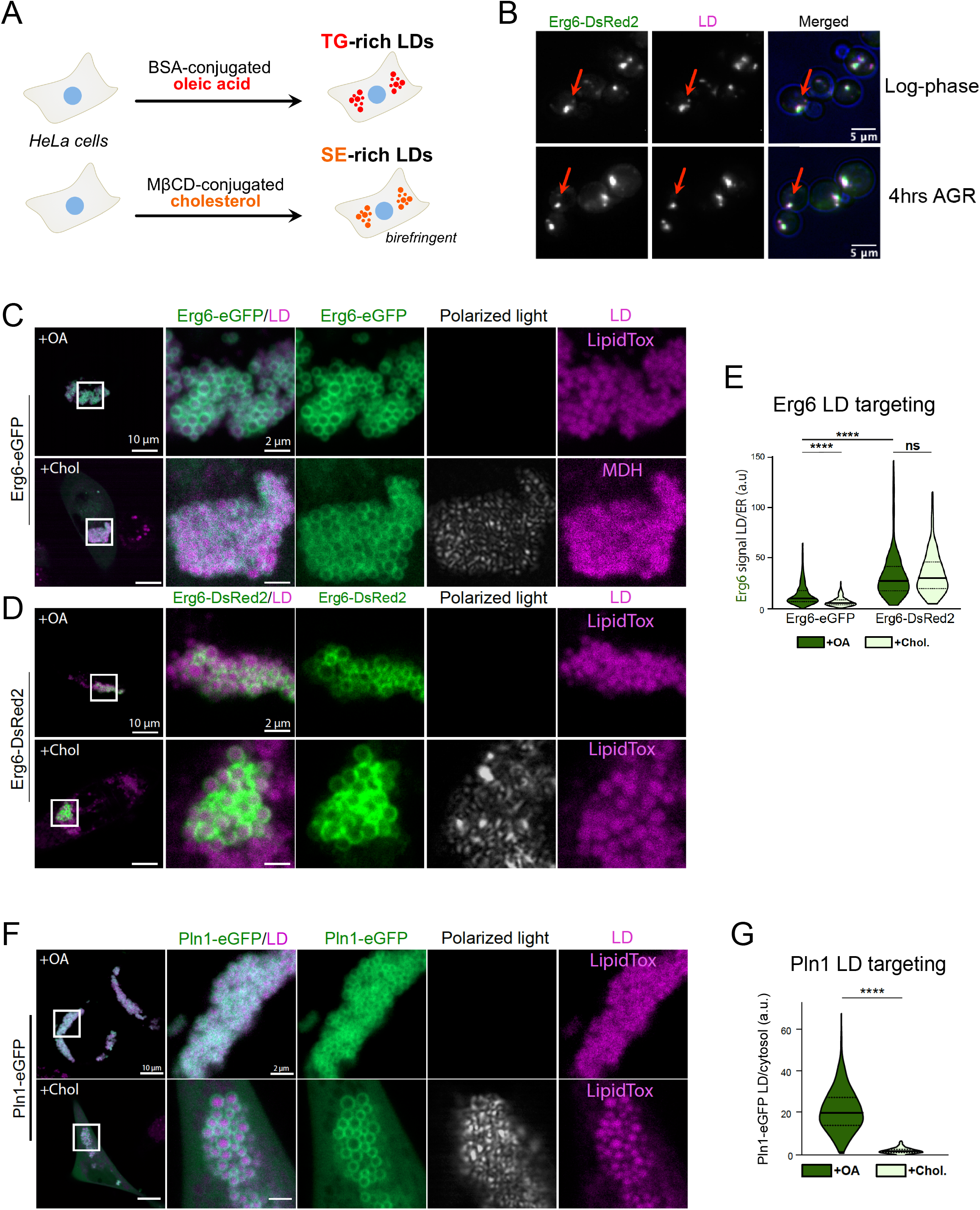
Erg6 and Pln1 exhibit LD targeting preferences in human HeLa cells independent of AGR stress. **A**) Cartoon of HeLa cell lipid treatments with BSA-conjugated oleic acid (OA) or methyl-β-cyclodextrin-conjugated cholesterol (Chol), generating TG-rich or SE-rich LDs. **B**) Yeast expressing Erg6-DsRed2 and stained for LDs with MDH, then imaged either in log-phase growth (top) or following 4hrs AGR. **C**) HeLa cells transfected for 24hrs with Erg6-eGFP and then incubated with 400μM of oleic acid (OA) or 250μM of methyl-β-cyclodextrin conjugated cholesterol for 24hrs. Scale bars indicated. LDs stained with LipidTox or MDH. **D**) HeLa cells transfected for 24hrs with Erg6-DsRed2 and then incubated with 400μM of OA or 250μM of methyl-β-cyclodextrin conjugated cholesterol for 24hrs. **E**) Analysis of the quantified LD-to-ER targeting ratios of Erg6-eGFP or Erg6-DsRed2 after induction of TG or SE-rich LDs. Mean ± SEM n = 340 to 989 LDs from 19 to 28 cells. Triplicate expts were conducted. **F**) HeLa cells transfected for 24hrs with Pln1-eGFP and then incubated with 400μM of oleic acid (OA) or 250μM of methyl-β-cyclodextrin conjugated cholesterol for 24hrs. Scale bars indicated. LDs stained with LipidTox. **G**) Analysis of the quantified LD/cytosol targeting ratios of Pln1-eGFP after induction of TG or SE-rich LDs. Mean ± SEM n = 19-21 cells. (n=893 for TG-rich LDs, 268 for SE-rich LDs from (respectively) 14 and 12 cells). Triplicate expts were conducted. All scale bars indicated.

Previous studies dissecting LD protein targeting indicate that protein homo-oligomerization promotes a stable protein association with the LD surface, and this multimerization is thought to give perilipin proteins long-term associations with LDs (Giménez-Andrés et al., 2021). To test if protein homo-oligomerization could enhance Erg6 targeting to LDs, we tagged Erg6 with DsRed2, a well-established tetrameric fluorescent tag that we previously characterized as enabling protein oligomerization *in vivo* (Rogers et al., 2021). Indeed, when expressed in yeast Erg6-DsRed2 now decorated LDs in both log-phase as well as 4hrs AGR-treated conditions and no longer marked the ER, suggesting that the Erg6-DsRed2 tag promoted its retention on LDs even during AGR (**Figure 4B**). Next, we expressed Erg6-DsRed2 in HeLa cells treated with either OA or cholesterol. Erg6-DsRed2 exhibited significant targeting to both TG-rich and SE-rich LDs, and its calculated LD-to-ER ratios were elevated compared to Erg6-eGFP, as well as now equivalent between the TG-rich and SE-rich conditions (**Figure 4D,E**). These observations support a model where Erg6 LD targeting is influenced by the neutral lipid content within LDs, but LD targeting may be enhanced by protein multimerization.

Next, we expressed eGFP-tagged Pln1 in human HeLa cells treated with either OA or cholesterol. Notably, Pln1-eGFP clearly targeted to the surfaces of LDs in both conditions, indicating it can associate with both TG-rich as well as SE-rich LDs exhibiting birefringent LCL SEs. However, a dim Pln1-eGFP cytosolic signal was detectable in SE-rich samples. In line with this, quantification of the LD-to-cytosol Pln1-eGFP LD targeting ratios revealed significantly more targeting to TG-rich LDs compared to the SE-rich LDs (**Figure 4F,G**). This is consistent with previous reports that Pln1 (Pet10) displays preference for TG-rich LDs in yeast (Gao et al., 2017). Collectively, this indicates that both Erg6 and Pln1 are capable of targeting to TG-rich as well as SE-rich LDs manifesting birefringent LCLs. However, both proteins exhibit reduced targeting to SE-rich LDs, suggesting the neutral lipid content within the LD core influences their degree of LD targeting. It supports a model where the changes in LD targeting observed in yeast are due primarily to alterations in LD neutral lipids, rather than other indirect metabolic changes associated with AGR stress. Although Pln1 displayed retention on LDs during yeast AGR, we speculate that other factors such as protein-protein interactions on the yeast LD surface may further influence this LD localization.

### Proteomics and imaging reveal that AGR selectively remodels the LD proteome

Given the different localization patterns of Erg6 and Pln1 to TG-rich and SE-rich yeast LDs, we next sought to comprehensively map changes to the yeast LD proteome upon AGR-induced LCL-LD formation. We used density gradient centrifugation to isolate LDs from yeast grown either in log-phase or AGR stress (**Figure 5A**). To evaluate the quality of our LD isolation protocol, we performed Western blotting of whole-cell lysates and the subsequent isolated LD fractions. We observed a clear de-enrichment of mitochondrial protein Por1 and the plasma membrane protein Pma1 in the LD fractions, and an enrichment of Pln1 in the LD fraction, suggesting this fraction was relatively pure (**Figure 5B**).

**Figure 5:**
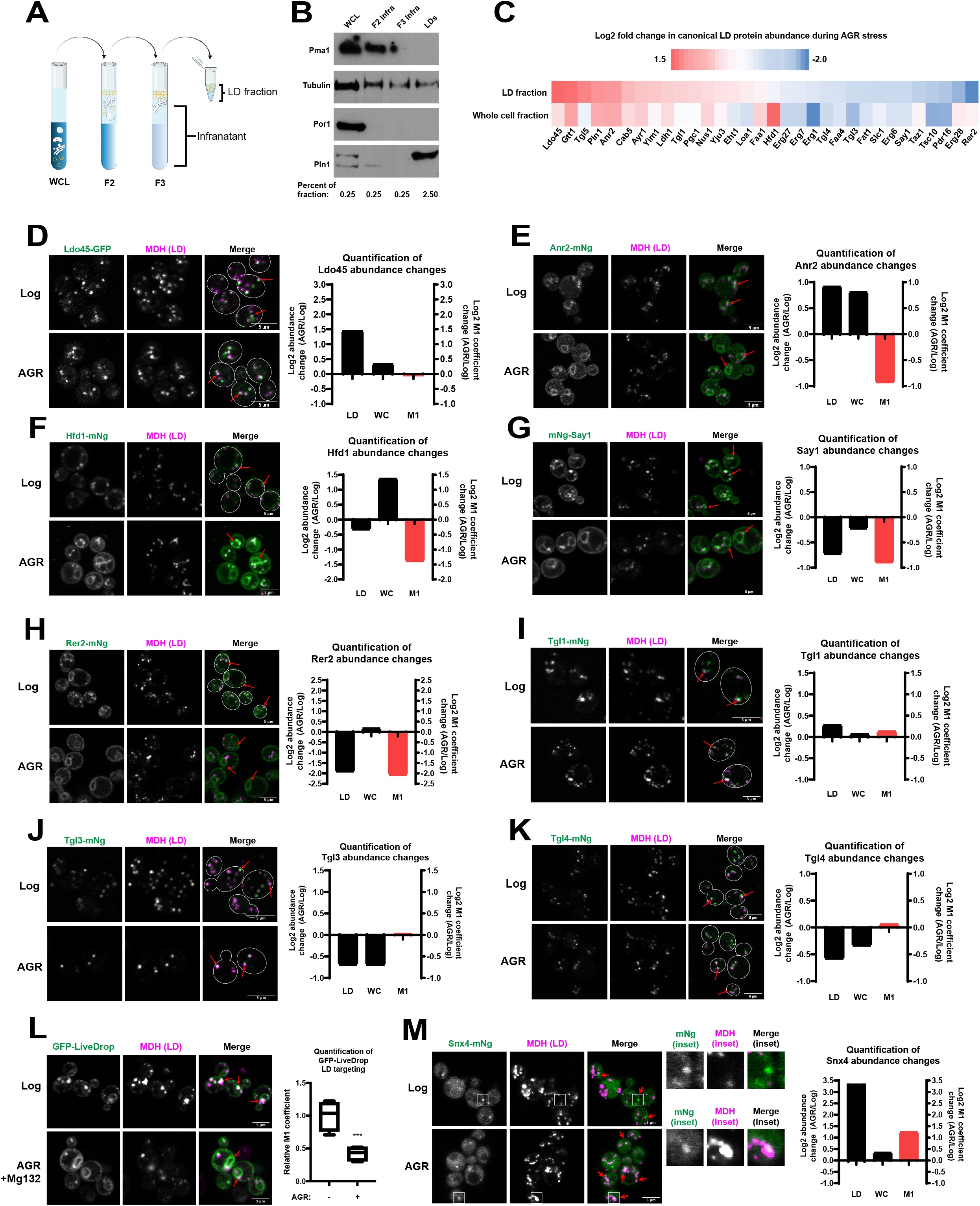
Proteomics and fluorescence imaging indicates selective remodeling of LD proteome during AGR. **A**) Cartoon schematic of step-wise LD isolation. **B**) Western blot of whole cell lysate (WCL), and fractions of LD isolation protocol as in A. Pma1: plasma membrane marker, Por1: mitochondria marker. Pln1: LD marker. Tubulin: cytoplasmic marker. **C**) Heatmap of Log2 fold abundance changes of annotated LD proteins from LC-MS/MS proteomics on either isolated LD samples (top) or whole-cell lysates (bottom). Heat map shows relative Log2 change of 4hrs AGR over log-phase samples. Yeast with GFP or mNeongreen (mNg)-tagged LD proteins with MDH LD stain in log-phase and 4hrs AGR. Each protein has quantification of log2 fold abundance changes from LD fraction (LD) and whole-cell (WC) proteomics as in panel C, as well as the relative M1 Manders coefficient from imaging. **L**) Yeast expressing GFP-Livedrop and imaged in either log-phase or 4hrs AGR conditions, with corresponding M1 Manders coefficient quantified. **M**) Snx4-mNg yeast with LD MDH stain in log and 4hrs AGR yeast. Insets are of LDs and Snx4-mNg signal. Calculated log2 fold abundance changes for LD, WC, and M1. All scale bars 5μm.

To establish a high-confidence LD proteome, we performed LC-MS/MS proteomics on both the floating LD fraction and lower infranatant fraction of log-phase and 4hrs AGR samples. Proteins were only considered LD-associated if they were: 1) more abundant in the LD fraction than the respective infranatant, and 2) detected in at least three of the four replicates for each growth condition. Of the 3188 proteins identified by LC-MS/MS, 167 proteins fit our criteria to be considered LD-associated. Within this dataset, we identified 32 of the 35 ‘canonical’ LD-associated proteins annotated in previous studies (Currie et al., 2014). Initially restricting our analysis to only these canonical LD proteins, we found an agreement between our yeast fluorescent imaging and proteomics. For example, Erg6 abundance was decreased in the proteomics of LD fractions taken from AGR yeast compared to log-phase, while Pln1 abundance increased in the proteomics of AGR yeast (**Figure 5C**, **top heat map row**). Remarkably, of the 32 LD proteins analyzed, some exhibited increased abundances in LD fractions from AGR yeast, while others had decreased abundances, suggesting significant proteome differences between the two samples. However, we considered that proteomics-based changes in protein abundance may not solely be attributed to changes in LD association, but could also reflect changes in whole-cell protein abundance. To distinguish between these possibilities, we also plotted the corresponding change in whole-cell abundance for each LD-associated protein during AGR (**Figure 5C**, **bottom heat map row**), as well as compared both the LD and whole-cell abundances directly on a 2-axis plot (**Figure S4A**). Indeed, this revealed that many canonical LD proteins also changed in their whole-cell abundances during AGR treatment.

To resolve this issue and better understand how protein localization to LDs was altered during yeast AGR, we fluorescently-tagged several canonical LD proteins and directly assessed their localizations using microscopy. Consistent with our proteomics data, Ldo45-GFP was localized to LDs during both log-phase and AGR stress conditions (**Figure 5D**). Also consistent with their proteomics profile, Hfd1-mNg, mNg-Say1, and Rer2-mNg targeted to LDs in log-phase yeast, but showed reduced (Hfd1, Say1) or even undetectable (Rer2) LD localization in AGR by imaging. Notably, these three proteins also exhibited elevated ER localization during AGR (**Figure 5F, G, H**). In this regard, Hfd1, Say1, and Rer2 behaved very similarly to Erg6-mNg localization, and exhibited similar decreases in Manders M1 coefficients in AGR stress that coincided with decreased abundances on isolated LDs by proteomics. However, imaging revealed some inconsistencies between the LD proteomics and fluorescence microscopy. For example, imaging indicated that Anr2-mNg was delocalized from LDs to the ER network during AGR, despite the LD proteomics suggesting its enhanced LD association in AGR stress (**Figure 5E**). This may be because Anr2 was also significantly more abundant in the whole-cell proteomics of AGR-treated yeast compared to log-phase, suggesting its LD abundance change may be due to its overall increase in cellular abundance.

Since TG lipases were required for LCL-LD formation in AGR (**Figure 1G, M**), we also monitored their sub-cellular localization by fluorescence microscopy. As expected, the TG lipases Tgl3-mNg, Tgl4-mNg, and Tgl5-mNg as well as Tgl1-mNg (a SE lipase) all decorated LDs in log-phase yeast (**Figure 5I,J,K, Figure S4B**). Remarkably, all four Tgl proteins retained LD localization following 4hrs AGR. This was generally in agreement with their LD proteomics profile, which suggested Tgl5 and Tgl1 had slightly elevated LD abundance during AGR stress (**Figure 5C**). While the LD proteomics suggested that Tgl3 and Tgl4 had decreased LD abundance in AGR, this may be due to their significant decreases in whole-cell abundances during AGR treatment (**Figure 5C**). It should be noted that the visual presence of TG lipases on LDs during AGR is consistent with a model where these lipases would be capable of the TG lipolysis necessary to alter the TG:SE neutral lipid ratios of LDs during the AGR time-course (**Figure 3B**), and thus promote TG:SE lipid demixing and lipid phase transitions.

Altogether, these data suggest that AGR stress and its associated LCL-LD formation promote the spatial re-distributions of many canonical LD-associated proteins, several of which de-enrich from LDs and re-target or accumulate at the ER network during AGR stress. However, a subset of proteins like perilipin Pln1 and Tgl lipases remain detectably LD-associated in yeast AGR.

### LiveDrop re-distributes from LDs to ER network during AGR

The observation that several LD resident proteins re-distributed in AGR treatment from LDs to the ER network was reminiscent of so-called Type I LD proteins, which move between the ER and LDs via lipidic bridges connecting the organelles (Wang et al., 2016). To interrogate whether Type I LD proteins could be re-targeted from LDs to the ER during LCL-LD formation, we monitored GFP-tagged LiveDrop, a model Type I minimal peptide from the LD protein *Dm*_GPAT4. As expected, GFP-LiveDrop localized predominantly to LDs in log-phase yeast, but a dim ER network signal was also detected, consistent with its dual organelle targeting (**Figure 5L**). However, following 4hrs AGR treatment GFP-LiveDrop was more prominently at the ER network, and LD association was significantly reduced when calculated with the M1 Manders coefficient (**Figure 5L**). This suggests that AGR-induced LCL-LD formation may promote the re-distribution of Type I LD proteins from LDs to the ER network.

### Snx4 associates with LDs during AGR stress

Thus far we have examined how AGR affects the localizations of canonical LD proteins. We also wanted to investigate whether proteins not typically associated with LDs may exhibit enhanced LD association during AGR stress. To this end, we queried whether such “non-LD” proteins were significantly elevated in the isolated LD proteomics of AGR-treated yeast. Indeed, the protein showing the greatest enrichment in these LD fractions was Snx4, a membrane binding sorting nexin typically associated with vesicle trafficking that can localize to endosomes and also autophagosomes (Nice et al., 2002), (Hettema et al., 2003) (**Figure S4A, right side of graph, Supplemental Table S1**). Notably, the Snx4 proteomics profile indicated it was significantly more abundant in isolated LD fractions from AGR yeast versus log-phase (horizontal plot), while its whole-cell abundance change between these conditions was mild (vertical plot), suggesting the enhanced LD proteomics abundance may be primarily due to *bone vide* increased LD targeting (**Figure S4A**, **Figure 5M graph**). We mNg-tagged Snx4 and evaluated its localization in log-phase and 4hrs AGR-treated yeast. As expected in log-phase, Snx4-mNg localized predominantly to the cytoplasm and punctate structures that did not overlap with LDs, which is consistent with its annotated localization to endomembrane compartments (**Figure 5M**). In contrast, at 4hrs AGR several Snx4-mNg foci co-localized with LDs, and the quantified Snx4-mNg Manders M1 coefficient exhibited an increase compared to log-phase, coinciding with its increased abundance in LD fractions by proteomics. Collectively, this suggests that during AGR, proteins not typically associated with LDs like Snx4 may display some LD targeting, possibly as a consequence to changes in the surface properties of the LD monolayer.

### Whole-cell proteomics suggests AGR promotes peroxisome fatty acid oxidation

Glucose restriction is a well-studied nutrient stress in yeast, and drives metabolic remodeling favoring the reorganization of organelles and utilization of alternative carbon sources when glucose is limiting (Seo et al., 2017), (Marini et al., 2020), (Eisenberg and Büttner, 2014). However, how glucose limitation alters lipid metabolism is underexplored. Since we conducted whole-cell LC-MS/MS proteomics of log-phase and 4hrs AGR yeast, we examined these datasets to determine whether changes in whole-cell protein abundances revealed patterns of metabolic remodeling that involved LDs and their lipids. We found that 4hrs AGR stress induced changes in the abundances of many proteins involved in fatty acid metabolism. In particular, peroxisome enzymes involved in fatty acid oxidation (FAO), including Pot1, Fox2, and Cta1 were among the most increased in abundance during AGR stress compared to log-phase growth (**Figure 6A, right side of plot**). Also elevated were the peroxisome-associated fatty acyl-CoA ligase Faa2 (which can promote import of fatty acids into peroxisomes), the acetyl-CoA transporter Crc1 (which transports acetyl-CoA derived from peroxisomal FAO to mitochondria), as well as Yat1, a carnitine acetyl-transferase that works with Crc1 to promote mitochondrial acetyl-CoA utilization. Enzymes related to the glyoxylate cycle including Icl1 and Idp2, the malate synthase Mls1, and acetyl-CoA synthase Acs1 were also among the most elevated proteins in AGR-treated yeast, suggesting pathways to generate and use acetyl-CoA pools were elevated in glucose restriction (**Figure 6A,B**). In contrast, amino acid transporters like Mup1 and Lyp1 were significantly decreased in abundance (**Figure 6A, left side of plot**), consistent with their turnover during glucose starvation that promotes adaptive metabolic remodeling (Lang et al., 2014), (Wood et al., 2020).

**Figure 6:**
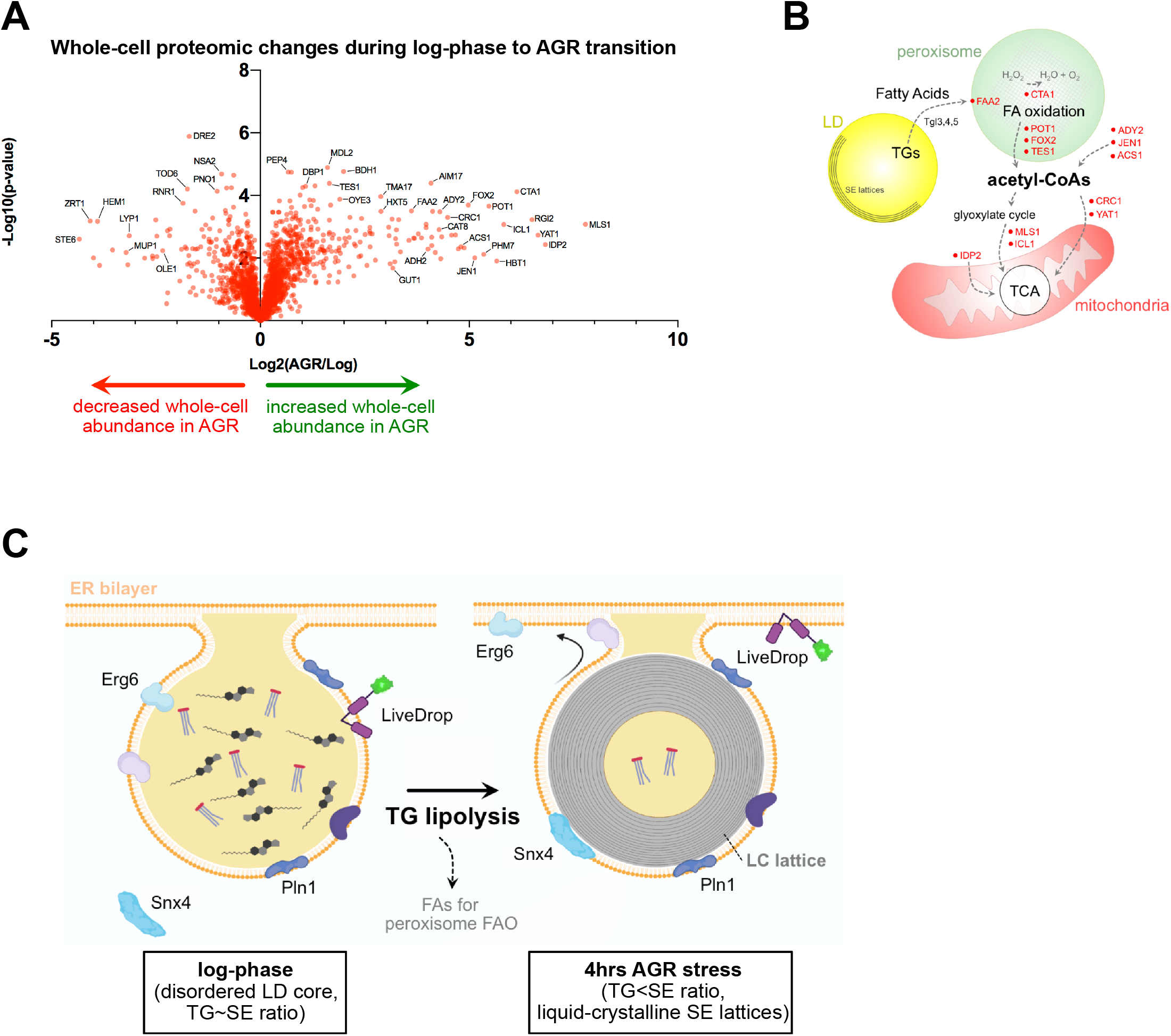
Glucose restriction promotes metabolic remodeling favoring peroxisome fatty acid oxidation. **A**) Volcano plot showing log_10_ p-value and log_2_ abundance changes in wholecell abundance of proteins in 4hrs AGR treatment versus 2% glucose log-phase growth. Proteins on right are increased in whole-cell abundance with 4hrs AGR, those of left decreased in abundance. Data are average of 4 independent expts. **B**) Schematic of inter-organelle interactions from proteomics depicted in panel A. **C**) Cartoon of working model for AGR-induced LCL-LD formation, and protein re-targeting of various LD proteins.

Collectively, this indicates that glucose restriction promotes the mobilization of TGs from LDs, providing fatty acids and ultimately acetyl-CoA as fuel for cellular energetics. Indeed, acetyl-CoA generated by peroxisome FAO can be delivered to mitochondria to fuel its energetics in the absence of glucose, suggesting inter-organelle remodeling during AGR that enables LD-derived lipids to fuel alternative carbon metabolism. In this model, an additional consequence of TG mobilization is a shift in the TG:SE neutral lipid ratios within LDs, ultimately giving rise to neutral lipid demixing and SE phase transitions into smectic liquid-crystalline phase lipids within LDs (**Figure 6C**).

## Discussion

Emerging evidence suggests that the phase transition properties of cellular biomolecules, such as proteins in membraneless organelles, directly influence organelle function and cell physiology. Like proteins, lipids can also undergo phase transitions. SEs can form liquid-crystalline lattices that are observed in human diseases like atherosclerosis, or in organelles like LDs. The metabolic cues that drive these phenomena, and their impact on organelle and cell physiology are unclear. Here we show that in yeast, glucose restriction promotes the formation of liquid-crystalline lattices within LDs. These lattices require TG lipolysis, and our experimental and modelling data support a model where TG is mobilized to support cellular energetics while simultaneously altering the TG:SE ratios within the LD hydrophobic core, promoting the spontaneous de-mixing of LD neutral lipids and the subsequent accumulation of SEs at the LD periphery beneath the surface monolayer. This lipid demixing would promote the transition of SEs from an amorphous to a smectic liquid-crystalline phase. Modelling also indicates that SE accumulation at the LD periphery alters the monolayer surface, creating shallow LPDs that could influence LD protein targeting. In line with this, we experimentally observe numerous changes to the LD proteome during AGR, suggesting changes to the LD core influence protein targeting to the LD surface, and protein spatial distribution between LDs and the ER network.

How proteins target LDs is still poorly understood, and can involve trafficking from the ER network or cytoplasm to the LD surface. In this study, we revealed that the LD proteome dramatically differs between AGR-treatment and log-phase growth. Erg6, a yeast canonical LD protein, relocalizes to or is retained at the ER network during AGR, suggesting it moves from LDs to the ER via lipidic bridges. This LD delocalization is suppressed in conditions that attenuated LCL-LD formation (i.e. OA treatment or loss of TG lipolysis). It can also be quickly reversed when cells are briefly heated to 40°C, (i.e. above the predicted melting temperature of smectic-phase SEs), suggesting direct movement of proteins between LDs and ER via ER-LD connections. We also observe similar Erg6 LD-to-ER re-targeting in human HeLa cells containing SE-rich birefringent LDs, supporting the model where Erg6 re-localization is due to changes in LD lipid content, rather than other changes in cellular metabolism associated with yeast AGR stress. Additionally, Type I LD peptide GFP-LiveDrop, which under log-phase conditions targets primarily to LDs, appears more ER localized during AGR. Collectively, this suggests that Type I LD proteins may favor ER localization versus the surfaces of LCL-LDs. This also indicates that many yeast LDs maintain connections to the ER network and thus exhibit the lipidic bridges necessary for this inter-organelle trafficking, consistent with earlier work (Jacquier et al., 2011). We also observe that some proteins display more LD retention during AGR stress. This may be attributed to protein-protein interactions or oligomeric properties of those proteins on the LD surface. In support of this, we find that artificially multimerizing Erg6 with a DsRed2 tag promotes its retention at LDs during AGR, implying that protein oligomerization enhances LD residency, as it has previously been observed for some perilipins (Kory et al., 2015), (Giménez-Andrés et al., 2021).

Whereas Erg6 delocalized from LDs during AGR, TG lipases Tgl3,4,5 remained LD bound. Although their LD anchoring mechanisms are not fully understood, this implies that LDs can mobilize TG during the AGR time-course, gradually altering the TG:SE ratio in a manner that supports SE phase transitions. In support of this, *in silico* modeling of LDs indicates that some TG remains accessible near the LD surface even as SEs accumulate there. Fatty acids derived from these TG pools are likely substrates for peroxisome FAO, of which several key enzymes are elevated during AGR stress. The acetyl-CoA produced from FAO could also fuel mitochondrial energetic pathways, several proteins of which are also elevated. LCL-LDs also exhibited de-localization of enzymes like Hfd1, Rer2, and Say1. It is possible the re-distribution of these and other enzymes may influence their activities, and therefore promote metabolic remodeling during glucose restriction. Indeed, several Erg pathway enzymes also appeared deenriched from LDs during AGR by proteomics, and Erg1 is more enzymatically active at the ER than on LDs (Leber et al., 1998).

Our proteomic and imaging analysis also revealed that LDs may become associated with non-LD proteins during AGR stress. This included the membrane trafficking protein Snx4, which colocalized with some LDs during AGR stress. As sorting nexin proteins can contain membrane binding/inserting modules, it is possible that Snx4 associates with LDs during AGR by inserting into its monolayer surface, but this requires further study. Since the LD surface is normally densely coated with proteins, it is also possible Snx4 and other proteins may associate with the LD surface as it is partially uncoated of canonical LD proteins during AGR stress. Notably, we also detected an enrichment of Rab proteins Ypt1, Ypt32, and Ypt32 in isolated LD proteomics from AGR-treated yeast (**Figure S4A**). In line with this, other studies reveal Rab proteins on LDs in specific conditions (Bersuker et al., 2018). It is tempting to speculate that the lipid moiety present on many Rab proteins may be attracted to the monolayer surface of LCL-LDs during AGR stress.

This study is a significant step toward understanding how metabolic cues influence the lipid phase transition properties of organelles, and ultimately organelle lipid and proteome composition and function. Future studies will interrogate whether such changes in the LD proteome influence metabolic remodeling that ultimately enable yeast to adapt to glucose shortage.

## Materials and Methods

Please see STAR Methods for a full description of the Methodology.

## Supporting information

STAR Methods

## Acknowledgement

We thank Jonathan Friedman, and members of the Henne and Nicastro labs for helpful insights during this study. We thank Daniel Stoddard for management of the UTSW electron microscope facilities and training, and Gang Fu for some data acquisition. The UT Southwestern CryoElectron Microscopy Facility is supported in part by the CPRIT Core Facility Support Award RP170644. We would also like to thank the UTSW proteomics and live cell imaging facilities for their assistance with data collection and analysis. Finally, we would like to thank Dr. Joel Goodman for the Pln1 antibody. W.M.H. is supported by funds from the Welch Foundation (I1873), the NIH NIGMS (GM119768), NIDDK (DK126887), Ara Parseghian Medical Research Fund, and the UT Southwestern Endowed Scholars Program. S.R. is supported in part by a NIH T32 training grant (5T32GM008297). L.G., E.R., and D.N. are supported by the Cancer Prevention and Research Institute of Texas grant RR140082 to D.N. S.V. and V.Z. acknowledges support from the Swiss National Science Foundation (PP00P3_194807) and from grants from the Swiss National Supercomputing Centre (CSCS) under project ID s980 and s1131. A.R.T. is supported by ANR-18-CE11-0012-01-MOBIL and ANR-CE11-0032-02-LIPRODYN. This research was supported in part by the computational resources provided by the BioHPC supercomputing facility located in the Lyda Hill Department of Bioinformatics, UT Southwestern Medical Center.

## Supplemental Figure Legends

**Figure S1:**
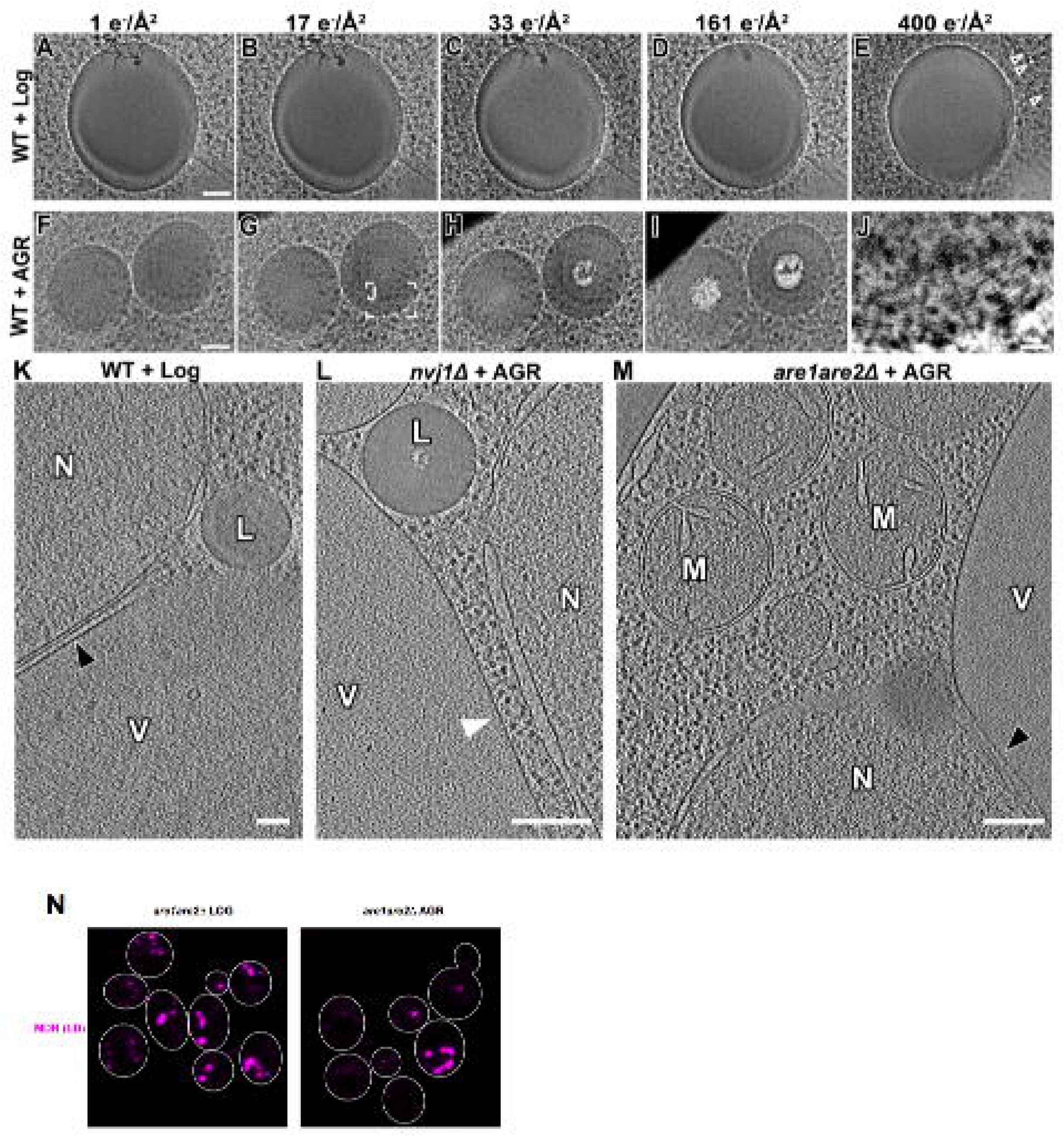
LD lipid phase transitions characterized by cryo-FIB and cryo-ET (corresponds to Figure 1). **A-J)** Electron dose series (“bubblegrams”) for LDs from cryo-FIB milled WT yeast in log phase (**A-E**) or after AGR (**F-J**); series of 2D cryo-EM images were recorded of the same LDs exposed to increasing electron dose (1 – 400 e-/Å2). Note that liquid-crystalline layers (LCLs) (see box in **G** magnified in **J**) and excessive bubbling in LD centers (starting at an electron dose <30 e-/Å^2^) occurred only under AGR. Even at 400 e-/Å^2^ electron dose, minimal bubbling (white arrowheads in **E**).was observed in log WT. **K-M)** Representative tomographic slices from cryo-FIB-milled and cryo-ET reconstructed WT in log phase (**K**), *nvj1Δ* yeasts after 4hrs AGR (**L**), and *are1are2Δ* yeast after 4hrs AGR (**M**). The nucleus-vacuole junction (black arrowheads in **K** and **M**) was observed in WT and *are1are2Δ* yeasts, but absent in *nvj1Δ* yeast (white arrowhead in **L**). No LDs were found in *are1are2Δ* yeast. V, vacuole. N, nucleus. L, lipid droplet, M, mitochondrion. **N**) Yeast stained with LD marker MDH in log and 4hrs AGR. Scale bars: 50nm (A-I), 200nm (K-M), 25nm (J).

**Figure S2:**
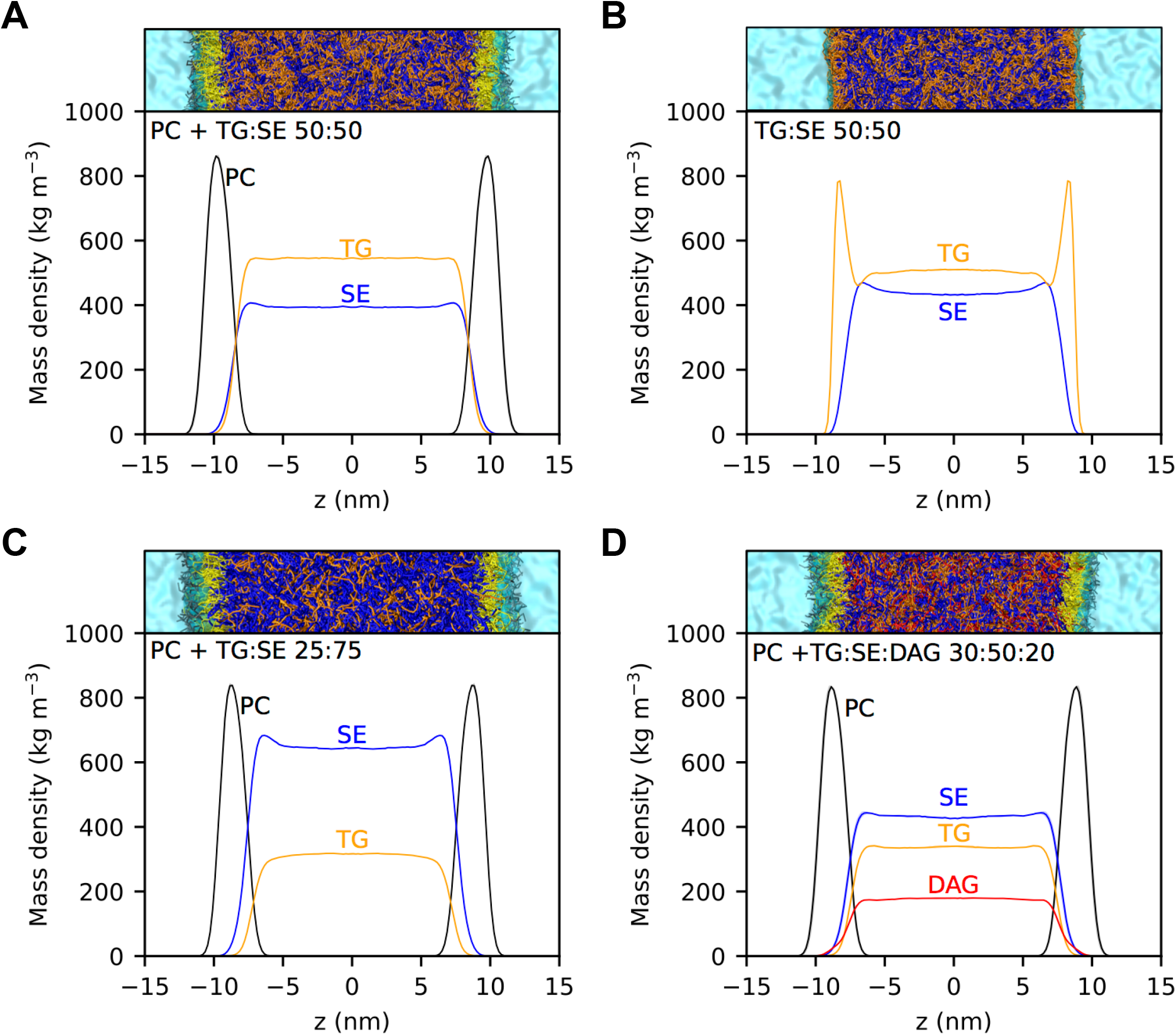
SEs and TGs partition non-uniformly in the core of LDs (corresponds to Figure 2). **A**) Density profile of TG, SE and PC in trilayer systems containing 50:50 TG:SE. For density profiles, the results are reported as the average between the analysis of the first part and the last part of the simulations, while the shaded regions correspond to the area between the two curves. See Methods for details. **B**) Density profile of TG and SE in systems containing 50:50 TG:SE with no PC phospholipids (PLs). **C**) Density profile of TG, SE and PC in trilayer systems containing 25:75 TG:SE. (**D**) Density profiles of PC, SE, TG and DAG in systems containing 30:50:20 TG:SE:DAG.

**Figure S3:**
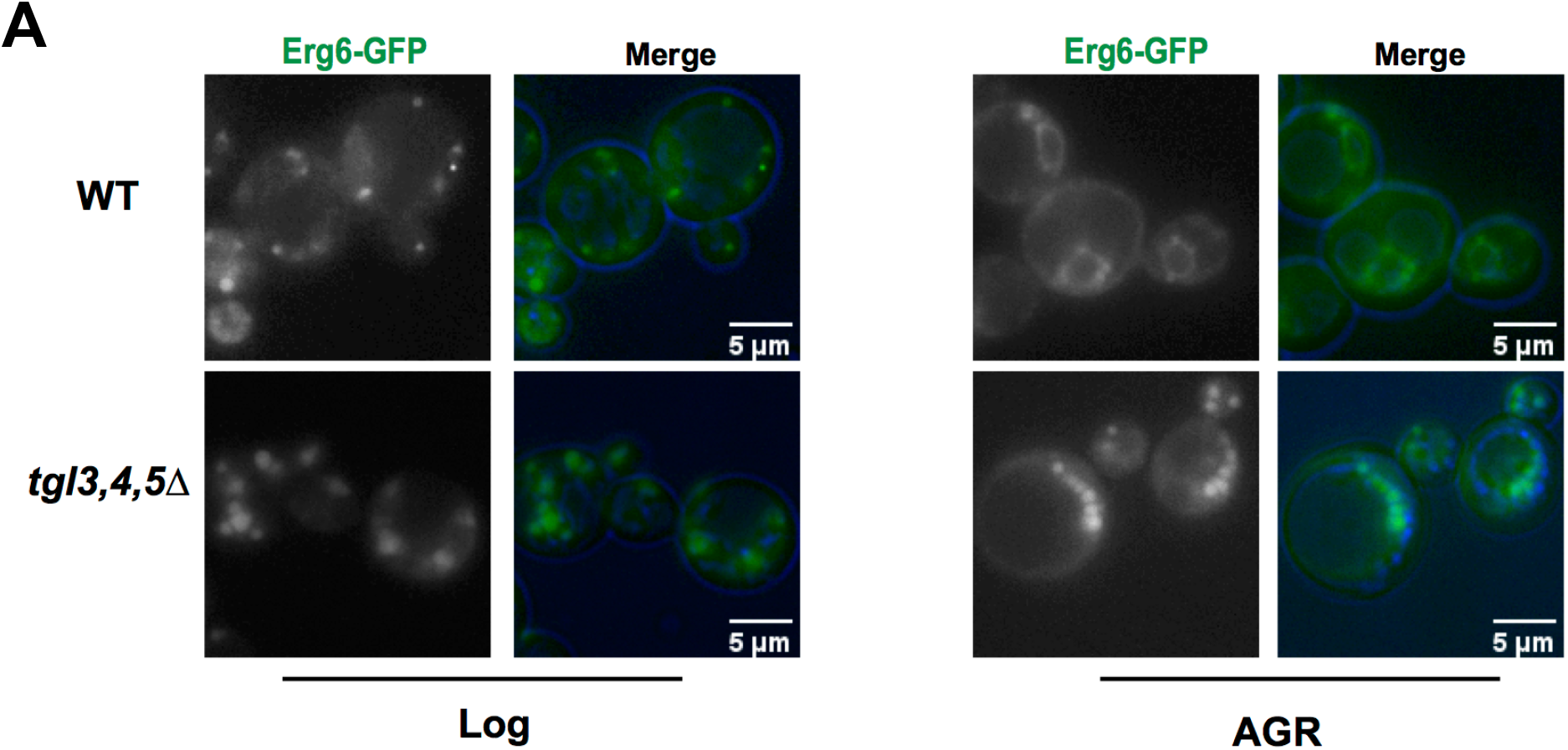
TG lipases are required for Erg6-GFP retargeting to the ER network during AGR-associated LCL-LD formation (corresponds to Figure 3). **A**) WT or *tgl3,4,5Δ* Erg6-GFP yeast in log or AGR conditions.

**Figure S4:**
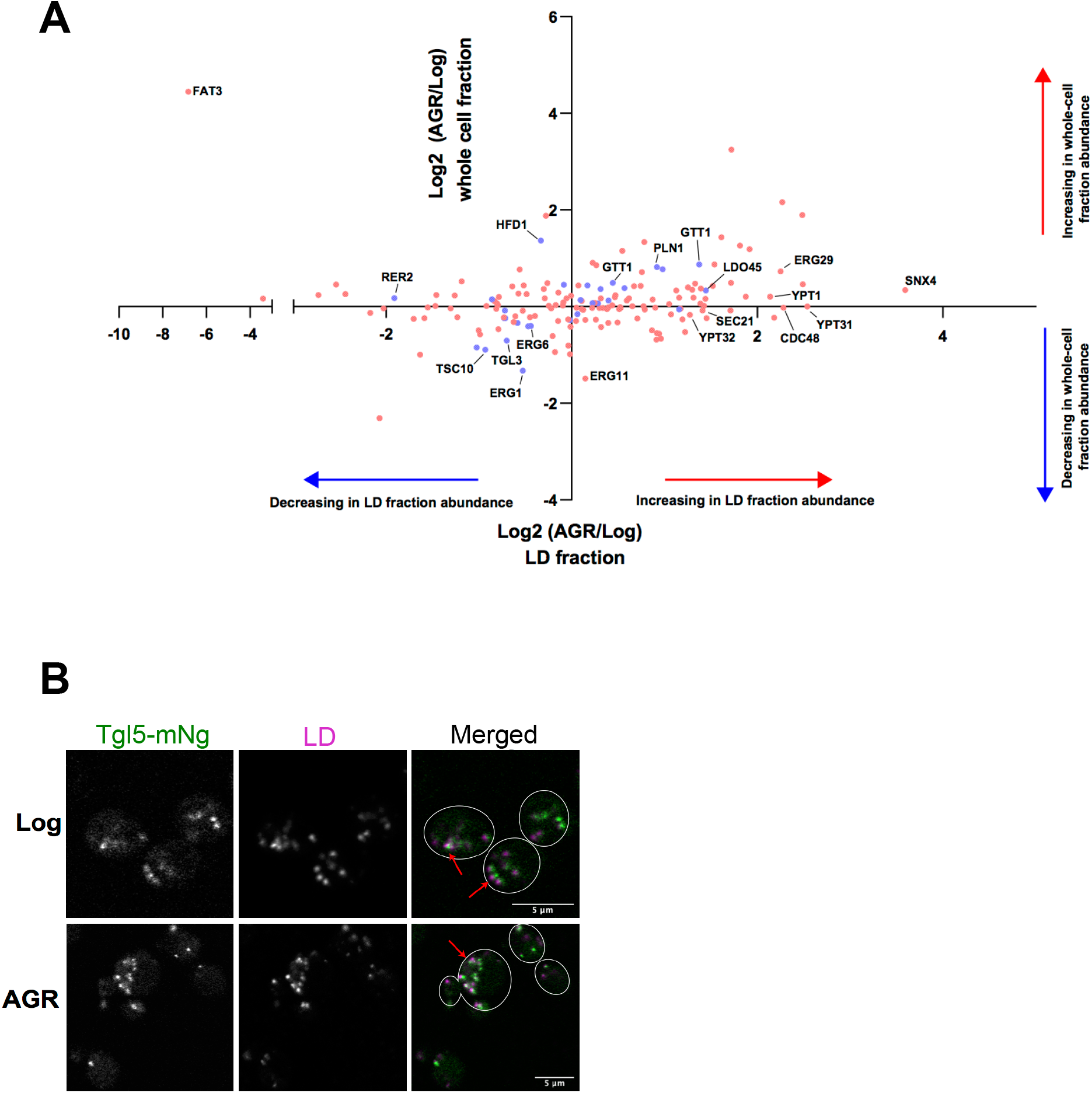
Selective re-localization of LD proteins during AGR stress (corresponds to Figure 5). **A**) Plot of protein abundances in whole-cell LC-MS/MS proteomics (vertical y-axis) versus their change in LD confidence score (horizontal x-axis). See methods for description of these values. Data are average of 4 independent expts. Blue indicate annotated LD proteins, and red indicate non-LD annotated proteins. **B**) Tgl5-mNg yeast co-stained for LDs with MDH. Scale bars 5μm.

**Supplemental Movie S1**: Tomographic reconstruction of a cryo-FIB-milled WT yeast cell after 4hrs acute glucose restriction. Compare with Figure 1A. Scale bar: 200nm.

**Supplemental Movie S2**: Tomographic reconstruction of a LD from a cryo-FIB-milled WT yeast cell in log phase (grown with 2% glucose). Compare with Figure 1B. Scale bar: 50nm.

**Supplemental Movie S3**: Tomographic reconstruction of a LD from a cryo-FIB-milled WT yeast cell after 4hrs AGR. Compare with Figure 1C. Scale bar: 50nm.

**Supplemental Table S1:** Spreadsheet of LD and whole-cell (WC) proteomics, with fraction analysis of 4 different log-phase and 4hrs AGR treated yeast samples. Columns A-I show the detected samples for the 8 independent experiments. Column W-Z is the ordered list of detected protei2ns, ordered with the most abundant proteins detected in LD proteomics from AGR-treated yeast (LCL-LD fractions) to least. Column AB is the corresponding change in WC abundance for each of these proteins. The No_WC_Data tab are proteins detected in the LD proteomics, but not detected in either one or both of the whole cell datasets.

